# Competing effects of indirect protection and clustering on the power of cluster-randomized controlled vaccine trials

**DOI:** 10.1101/191163

**Authors:** Matt D.T. Hitchings, Marc Lipsitch, Rui Wang, Steven E. Bellan

## Abstract

Power considerations for trials evaluating vaccines against infectious diseases are complicated by indirect protective effects of vaccination. While cluster-randomized trials (cRCTs) are less statistically efficient than individually randomized trials (iRCT), a cRCT’s ability to measure direct and indirect vaccine effects may mitigate the loss of efficiency due to clustering. Within cRCTs, the number and size of clusters affects three determinants of power: the effect size being measured, disease incidence, and intra-cluster correlation. We simulate trials conducted in a collection of small communities to assess how indirect protection and clustering affect the power of cRCTs and iRCTs during an emerging epidemic. Across diverse parameters, we find that within the same trial population, cRCTs are never more powerful than iRCTs, although the difference can be small. We also identify two effects that attenuate the loss of cRCT power traditionally associated with increased cluster size. First, if enrollment of fewer, larger clusters is performed to achieve higher vaccine coverage within vaccinated communities, this increases the effect to be measured and, consequently, power. Second, the greater rate of imported transmission in larger communities may increase the attack rate and similarly mitigate loss of power relative to a trial in many, smaller communities.

---

Cluster-randomized controlled trials (cRCTs) have become an increasingly common method for evaluating interventions for infectious diseases, including vaccines. Compared to individually randomized controlled trials (iRCTs), cRCTs may offer logistical, operational, and acceptability advantages (1), and allow the measurement of direct and indirect effects of vaccination, which are often relevant for policy-makers (2). The statistical theory of cRCT design has largely focused on the effect of clustering, commonly measured by intracluster correlation, on power (3-5). Intracluster correlation arises because outcomes of members of the same cluster are more similar than those from different clusters. Therefore, increasing the number of individuals within a cluster provides less information than would adding the same number of individuals in a new cluster.

When the trial outcome is an infectious disease, correlation arises also because each case in a cluster can transmit infection to other cluster members. Thus, trials of vaccines against infectious diseases exhibit a more complicated relationship between statistical power and sample size than in trials for non-infectious outcomes (6, 7); in particular, the total or overall vaccine effect measured by a cRCT is generally larger than the direct effect measured by an iRCT. In principle, this increased effect size in a cRCT might partially or fully offset the loss of power due to within-cluster correlation. Understanding these complexities can aid in vaccine trial design for emerging epidemics. While an important consideration in any clinical trial, maximizing efficiency is particularly crucial in trials during infectious disease emergencies such as the 2014-16 Ebola epidemic, where evaluation of experimental vaccines is especially urgent, and where limited available vaccine doses and/or changing disease incidence may constrain trial design (8).

In this paper, we first compare the power of an iRCT with that of a cRCT in the same population across a broad range of realistic parameters, taking into account that the cRCT is generally measuring a larger effect size. We hypothesized that, when R_0_ is slightly above 1, a cRCT may have greater power to detect total vaccine effects than an iRCT would have to detect direct effects. Our justification was twofold. First, a vaccine’s total effect is greater than its direct effect and thus more easily detected. Second, when an iRCT is conducted within numerous small communities, the indirect effects of vaccination may reduce incidence amongst control participants sufficiently as to erode the trial’s power (6). In a second analysis, we restrict our attention to cRCTs and consider two decisions an investigator must navigate when balancing the number of clusters with the size of a cluster, for a given trial population size (Fig. 1). Throughout we distinguish between *communities* that are targeted for enrollment, and *clusters* that comprise the individuals enrolled. When study clusters are sampled from communities, the first decision (*enrollment proportion*) concerns whether to enroll a larger proportion of each community from fewer communities, or to enroll a smaller proportion from a larger number of communities, fixing community size. The second decision (*community size*) concerns whether to recruit clusters from a smaller number of large communities or recruit from a larger number of small communities, fixing enrollment proportion.

**Fig. 1.**
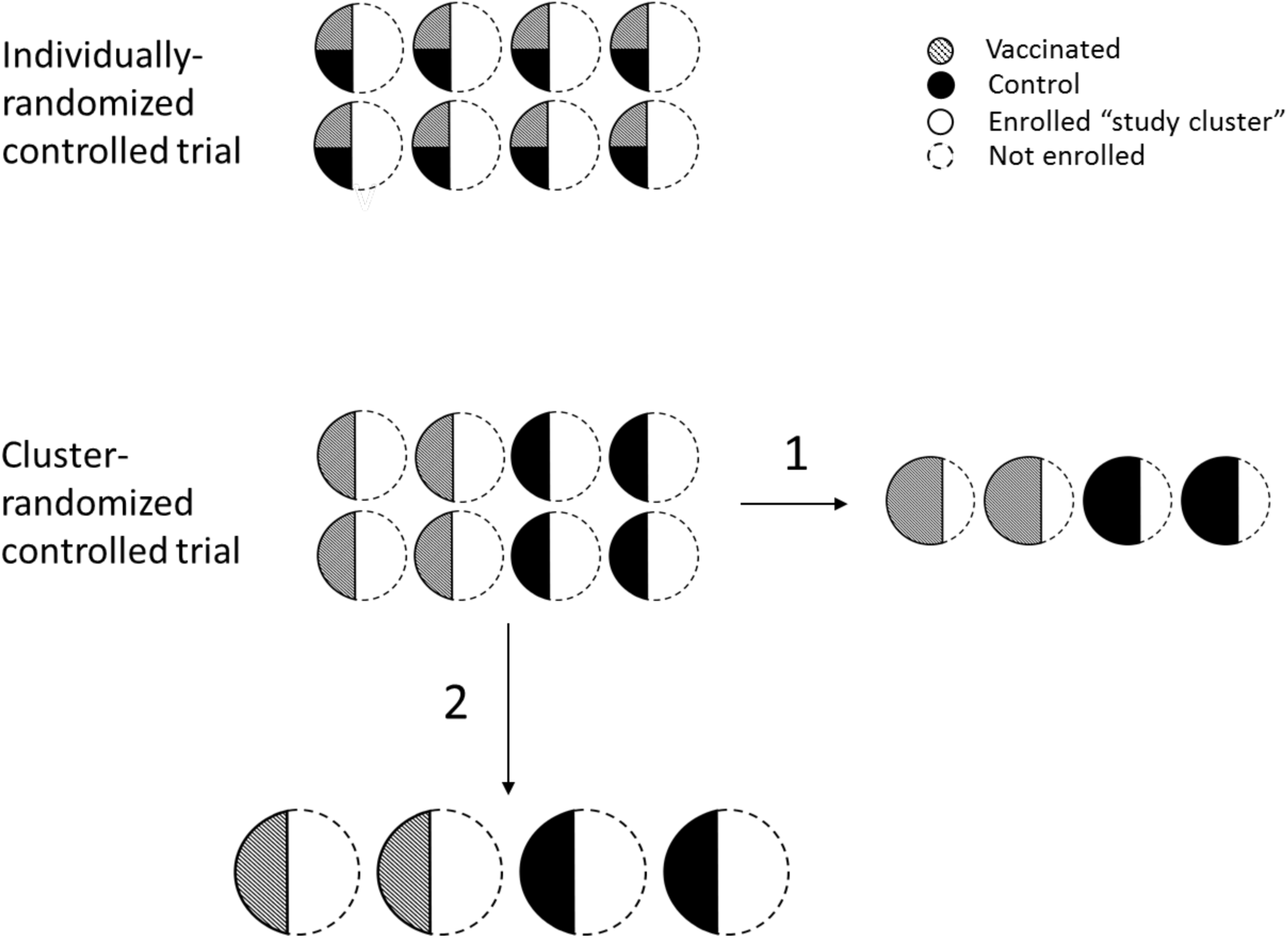
Schematic of an iRCT (top) and a cRCT (bottom). Study *clusters* (solid outlined) are enrolled from *communities* (circles). In the iRCT, individuals within each cluster are randomized to vaccine (striped) or control (black). In the cRCT, half the clusters are randomized to vaccine and half to control. In the cRCT design, fixing the number of individuals enrolled, there are two ways to balance cluster size and number of clusters in the trial: (**1**) fixing the community size, vary the enrollment proportion and the number of communities enrolled, and (**2**) fixing the enrollment proportion, vary the community size and number of communities.

With regard to enrollment proportion, recruiting a higher proportion of each community leads to higher vaccine coverage in communities receiving vaccination and thus more indirect protection to individuals therein. The greater overall protection may lead to increased power. With regard to community size, larger communities may experience an increased rate of introduction into the community if, for example, disease importations are proportional to the number of travelers to and from the community, which likely scales with community size. Both the increased indirect protection and the increased importation rate may increase power because they increase the effect size and the average number of cases in the trial population, respectively. These effects may thus partially counterbalance the loss of power that is known to accompany having fewer, larger clusters. We use a transmission model of an emerging directly transmitted infection (such as Ebola virus disease) to assess the contribution of these effects to the relative power of iRCTs and cRCTs.

## METHODS

### Theoretical Analysis

We first explored the plausibility that cRCTs may be more efficient than iRCTs by using theoretical final size equations to calculate the expected outbreak probability and attack rate in clusters, varying enrollment proportion, R_0_, and vaccine efficacy (see Supplementary Material for details). While this analysis provides some insight into the trade-off between indirect effects and clustering, we conducted the following simulation-based analyses to more realistically account for how epidemic stochasticity may increase variability between communities.

### Simulated population structure

We consider a population divided into two distinct groups: a main population in which a major epidemic is progressing, and a smaller population made up of multiple small *communities* from which the trial population is enrolled. The communities are represented with a stochastic block network model (9), in which contacts between individuals within the same block are far more common than those between blocks. This assumption is essential as it increases the strength of indirect effects within clusters relative to scenarios in which there is more between-cluster transmission (10). A connection between individuals in the network represents a single infectious contact per day, and we assume that the number of contacts per individual (degree) is Poisson-distributed.

### Transmission models

To balance realism with computational feasibility we rely on distinct transmission models for the main population and for the communities, using a deterministic compartmental model and a stochastic compartmental model respectively.

Both models use a susceptible-exposed-infectious-removed (SEIR) compartmental structure. We assume that infections are introduced into communities via transmission from the main population, and the daily hazard of infection for an individual is proportional to the prevalence of infection in the main population. The community-level rate of disease importation (“importation rate”) is defined as the number of cases per year arising solely as a function of these external transmission events. We assume that the importation rate varies with the size of the community. In particular, larger communities experience more disease importation events, with community importation rate *M_i_* increasing with 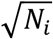, where *N_i_* is the size of the *i^th^* community (11). See Supplementary Material for more details on importation rate and disease natural history.

### Vaccine trial design

For both designs, the specified number of communities are enrolled on a fixed calendar day with a target proportion of community members enrolled at random from the susceptible and exposed individuals therein, forming that community’s *study cluster*. In the iRCT, half the individuals in each study cluster are randomized to vaccination with the other half to placebo control. In the cRCT all individuals in half the study clusters are assigned to vaccination, while those in the other half are assigned to placebo control. In this design, all enrolled individuals in clusters assigned to vaccination are vaccinated.

### Statistical analysis

Statistical analysis of the trial is based on time to symptom onset, with individuals censored after a fixed time. For the iRCT, a Cox proportional hazards (PH) analysis is performed to estimate the direct effect of the vaccine, stratifying by community (12). We define statistical significance at the α=5% level using a two-tailed Wald test, and for each combination of parameters we simulate 500 trials, estimating the power as the proportion of trials that reject the null hypothesis of no vaccine effect, which accounts for different estimands used by different designs. We calculate the median vaccine effect estimate across the simulated vaccine effect estimates. To estimate the Type I error of each design we repeat the above process with the true vaccine efficacy set to 0. To measure the magnitude of clustering in the cRCT we report the design effect, defined as *DE* = 1 + *ρ*(*m* − 1), where *ρ* is the intracluster correlation coefficient (ICC) calculated using (13), which is likely an underestimate of the ICC for time-to-event data (14), and *m* is the average size of a study cluster. The design effect increases with ICC, as subjects in the same cluster are more similar, and with the size of each cluster, as there are fewer, larger groups of similar individuals. The ICC is a measure of between-cluster variance relative to total variance in the outcome: if between-cluster variance is large relative to within-cluster variance, the ICC is large and individuals in the same cluster provide little information relative to individuals in different clusters.

In this cRCT design, a Cox PH model estimates the total effect of vaccination. To ensure we used a cRCT analysis that maintains nominal Type I error when comparing cRCT power to that of an iRCT, we first compared Type I error between several methods to account for clustering when determining statistical significance within the cRCT design: namely, a Cox PH model with Gaussian- or gamma-distributed shared frailty, and a Cox PH model with robust standard error estimate. We excluded from analysis individuals who developed symptoms within 10 days after vaccination (the average incubation/latent period) to avoid diluting the vaccine effect by analyzing infections that preceded vaccination.

### Choice of parameters

Table 1 shows the parameters used in the model, their meanings, values under baseline assumptions, range explored (where applicable), and references or justifications.

**Table 1.** Model Parameter Names, Values and Ranges Varied Across, Meanings and References or Justifications.

| Parameter | Meaning | Value/Range | Reference |
| --- | --- | --- | --- |
| $R_0$ | Average number of secondary infections generated by an infected individual | 0.6-3 | Wide range spanning most emerging infectious diseases. Calculated for network models using (22). |
| Mean (latent) | Mean latent period length (days) | 9.7 | (23) |
| SD (latent) | Standard deviation of latent period length (days) | 5.5 | (23) |
| Mean (infectious) | Mean infectious period length (days) | 5.0 | Time to hospitalization (23). |
| SD (infectious) | Standard deviation of infectious period length (days) | 4.7 | (23) |
| VE | Individual vaccine efficacy | 0.6 (0.4-0.8) | Baseline assumption. |
| $N_i$ | Size of community $i$ | 100 (50-200) | Assumption that some unit of this size exists in the population. |
| $M_i$ | Importation rate into communities | 0.025<br>(0.0125-<br>$0.05)\sqrt{N_i}$<br>cases/1,000<br>person-years | Based on a calculation for<br>measles (11), with the<br>magnitude of the rate chosen<br>so that there is on average<br>0.5 importations into a<br>community of size 100 over a<br>two-year epidemic. |
| Within-community<br>degree | Average total number of contacts<br>of an individual within the same<br>community | 14.85<br>(14.83-14.85) | Based on *Ebola, ça suffit!* trial<br>(20) (ring size of 90, <20% of<br>which were primary<br>contacts). |
| Between-<br>community<br>degree | Average total number of contacts<br>of an individual from outside their<br>community | 0 (0-0.02) | Assumption that communities<br>disconnected to minimize<br>spillover effect. A range was<br>explored to represent one or<br>two contacts outside each<br>community. |
| Trial size | Average number of individuals<br>enrolled | 4,000 | Assumption to achieve<br>reasonable power for chosen<br>parameters. |
| Trial start day | First day of enrollment, vaccination and start of follow-up, relative to the first day of the epidemic in the main population | 150 (100-250) | Assumed the trial starts before the peak of the epidemic in the main population and that the trial team is ready to go when epidemic starts. |
| Trial length | Length of follow-up after trial start (days) | 140 (70-210) | Assumption to achieve reasonable power for chosen parameters. |

## RESULTS

### Comparison of iRCT and cRCT

In our theoretical analysis based on final size calculations, we found support for our initial hypothesis that cRCTs could be more efficient than iRCTs: when R_0_ in vaccinated clusters in the cRCT is just above 1, the measured total effect is close to 1, which increases power; on the other hand, indirect effects in the iRCT drive down the incidence of disease among controls, undermining its power. Increasing enrollment proportion increases power of the cRCT relative to the iRCT, and there were parameter ranges for which the cRCT was more powerful than the iRCT; for example, with communities of size 100 and enrollment proportion 60%, we estimated that a cRCT would be more efficient than an iRCT when R_0_ was close to 1.6 and vaccine efficacy was between 50% and 60% (see Figure S2).

However, our simulation model reveals that, across a broad range of parameters, including population structure, trial design and vaccine efficacy parameters, iRCTs were always were more powerful than cRCTs in the same population, despite the larger effect size being measured in cRCTs. The discrepancy between the models arises because theoretical calculations underestimate the average cumulative incidence, as well as the variability in transmission across clusters, when R_0_ is close to 1. Figure 2 illustrates the power of simulated iRCT and cRCT designs versus R0, and highlights two findings. Firstly, the cRCT generally yields greater effect size estimates than the iRCT, because it measures the total vaccine effect rather than solely direct effects (Fig. 2A). Secondly, the design effect is large and increases with increasing R_0_ (Fig. 2B), because large R_0_ leads to more outbreaks within communities, which increases between-cluster variance and thus the ICC (see Fig. S1). Therefore, the power that the cRCT gains by measuring a larger effect is more than compensated by loss of efficiency due to within-cluster correlation. These two points explain why cRCT power first increases and then decreases with increasing R_0_. As R_0_ increases past a certain threshold, the effect of clustering begins to dominate the effect of increased incidence in the study population, and the trial loses rather than gains power from the increased transmission.

**Fig. 2.**
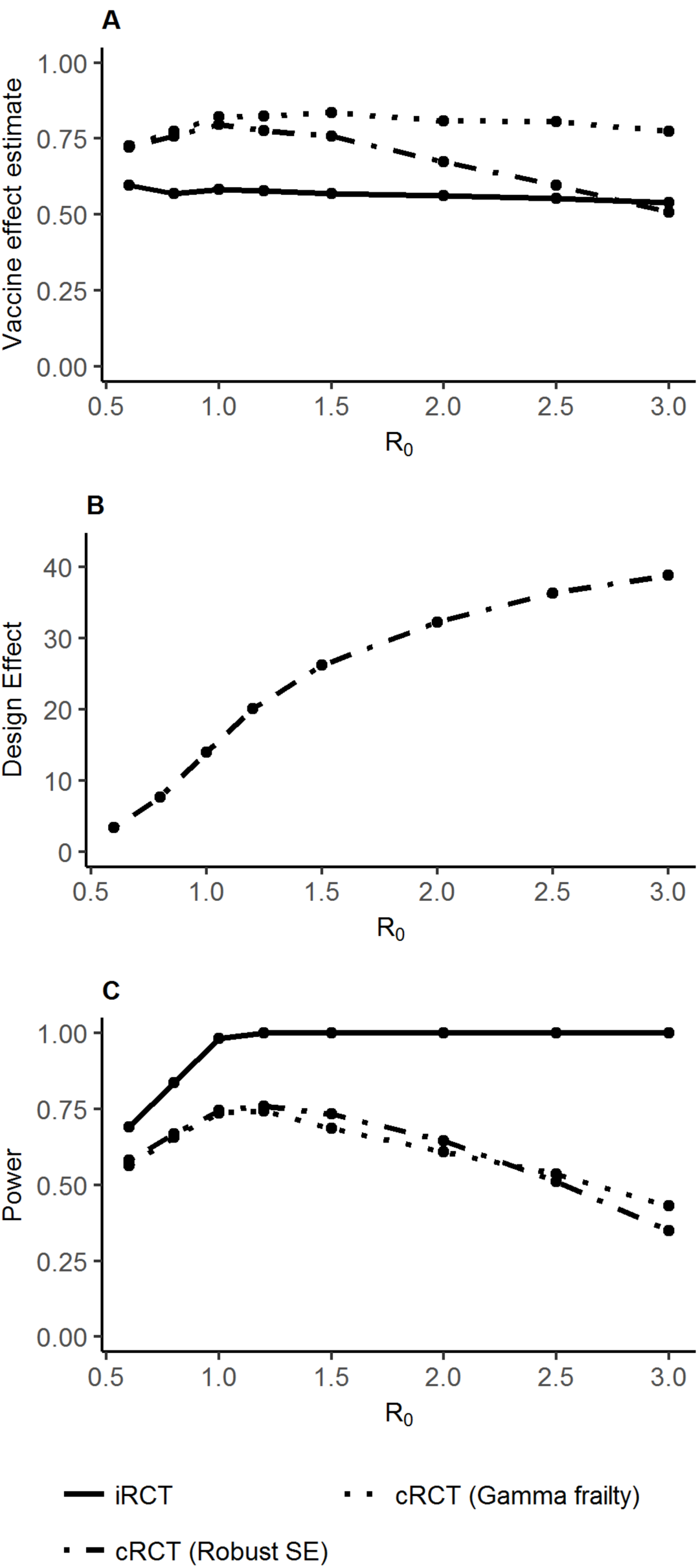
Comparison of vaccine effect estimates and power of individually- and cluster-randomized controlled trials. Vaccine effect estimates (**A**), design effect (**B**), and power (**C**) from an individually-randomized controlled trial (iRCT) and from a cluster-randomized controlled trial (cRCT) analyzed either using a shared gamma frailty model or using a Cox PH model with robust standard error estimates (robust SE). The incidence rate of importations into an average community is 0.5 cases/year, the vaccine efficacy is 60%, and other parameters are the baseline values listed in Table 1.

As hypothesized, we found that there was reduced incidence among controls in the iRCT compared to those in the cRCT due to indirect protection from vaccinated individuals (see online Shiny App at https://matthitchings.shinyapps.io/shiny), although this did not significantly affect the power of iRCTs in our simulations. This is likely because vaccine coverage was low in the iRCT (a maximum of 50% of individuals within clusters are vaccinated) such that there is still sufficient transmission amongst control participants to evaluate the vaccine, in part because importation events from the main population occur even in the presence of herd immunity.

The above results focus on the gamma-frailty model for analyzing the cRCT. We found that the estimated vaccine effect from a Cox PH model with robust standard errors (robust SE model) decreased drastically as R_0_ increases. This occurred because the effect estimate from the robust SE model is not stratified by cluster, and is thus biased by heterogeneity in hazard of infection caused by stochastic variation in outbreak size (12). The gamma-frailty model can account for this heterogeneity and performed better, yielding both Type I error rates below 5% and unbiased estimates of total vaccine effects for many of the parameter combinations. Still, when R_0_ was sufficiently small, the gamma-frailty model of cRCT designs did exhibit slightly elevated Type I error (see online <u>Shiny App</u>) due to sporadic and heterogeneous nature of outbreaks in the communities.

Figure 2 shows that the power of the cRCT is strongly affected by the design effect (Fig. 2C), and that the difference in power between the cRCT and iRCT is smaller when there is low R_0_. This observation held when other parameters were varied, including trial start day (relative to epidemic onset), vaccine efficacy, importation rate, and population structure. In the setting of low R_0_, epidemics will die out stochastically in most clusters experiencing one or more case importation. The cluster-level attack rates are thus close to zero and the between-cluster variance is small (see online <u>Shiny App</u>).

### Varying community enrollment proportion in a cRCT

Restricting attention to cRCTs, Figure 3 displays the vaccine effect estimate (Fig. 3A), design effect (Fig. 3B), and power (Fig. 3C) for a cRCT across varying *community enrollment proportions* (holding community sizes constant, but varying number of communities). As expected the estimate of total vaccine effect increases with increasing proportion enrolled because it increases vaccine coverage and, consequently, the indirect effects in vaccinated clusters. However, the increased effect size is counterbalanced by increases in the design effect (driven by larger clusters). Thus, for all values of R_0_ displayed except the highest considered R_0_=3 there is no clear trend in power with the community enrollment proportion. For R_0_=3, the simulations follow the trend generally expected for cRCTs in which the use of more, smaller clusters increases trial power.

**Fig. 3.**
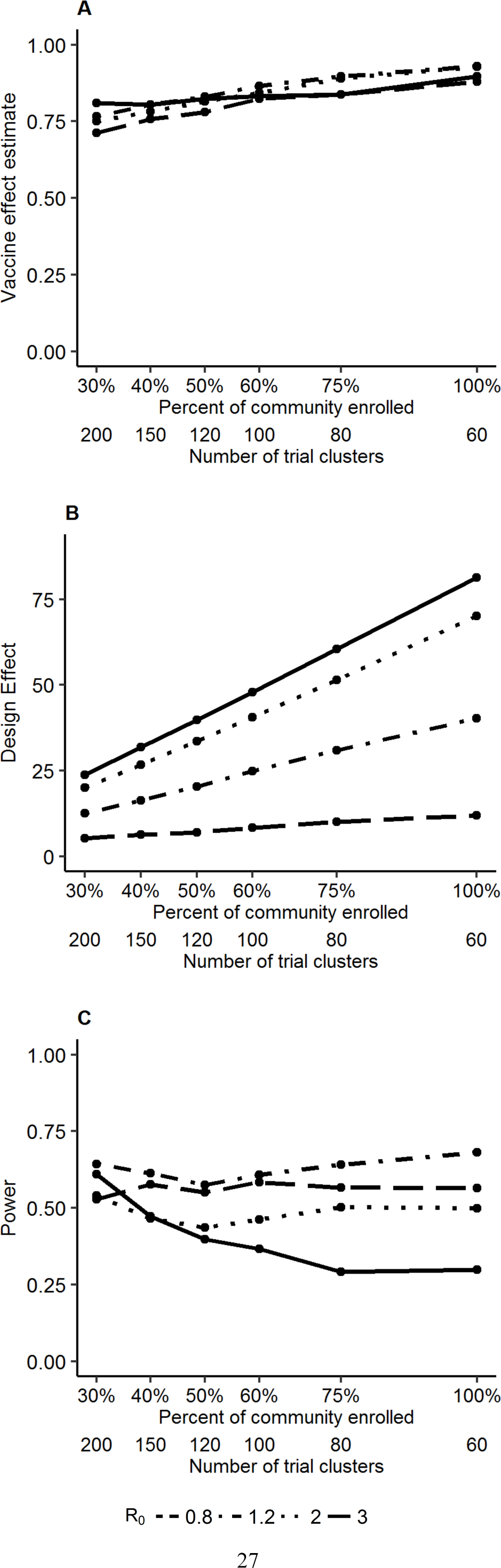
Relationship between power and community enrollment proportion for a cRCT. Vaccine effect estimates (**A**), design effect (**B**), and power (**C**) from a cRCT versus the percentage of individuals enrolled from each community, with total sample size held constant and assuming a vaccine efficacy of 60%.

### Varying size of enrolled communities in a cRCT

Figure 4 displays the attack rate in the study population (Fig. 4A), design effect (Fig. 4B) and power (Fig. 4C) for a cRCT with varying *size of enrolled communities*, holding the proportion of communities enrolled and the total number of trial participants constant. These results assume community importation rate scales with the square root of community size. As the community size increases, the per-community importation rate increases and with it the expected proportion of communities that receive an importation. When R_0_ is greater than 1 and many importations lead to outbreaks, fewer communities means that a higher proportion of communities experience outbreaks. In effect, to get fewer, larger communities one can imagine linking up small communities, meaning that an outbreak in one can spread to other small communities, boosting the attack rate. On the other hand, having many small communities means that outbreaks are limited by the size of the communities and the overall attack rate is lower. However, in this case the increased attack rate with fewer, larger communities is not large enough to offset the greater design effect and thus power decreases when increasing community size and decreasing community number.

**Fig. 4.**
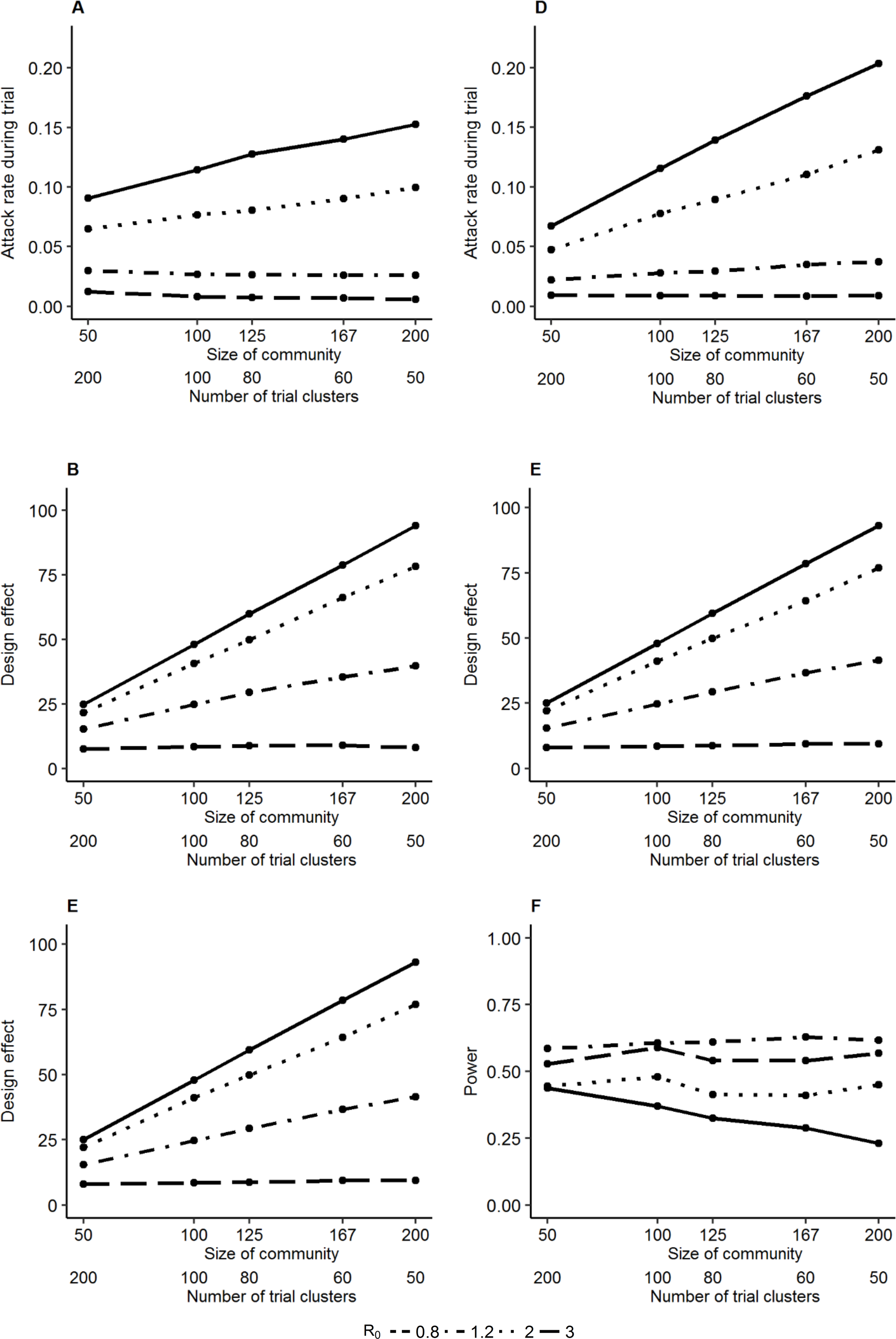
Relationship between power and size of enrolled communities for a cRCT. Attack rates in the trial population (**A** and **D**), design effects (**B** and **E**), and power (**C** and **F**) for cluster-randomized vaccine efficacy trials versus the size of the communities recruited, with total sample size held constant. In the left-hand column, community case importation rate is proportional to the square root of community size, and in the right-hand column it is proportional to the community size. All results shown here assume 60% community enrollment.

If community importation rate scales linearly with community size, there is an even greater increase in attack rate when there are fewer, larger communities (Fig. 4D), relative to the analysis above. In this case, even though the design effect increases with community size (Fig. 4E), the higher attack rate offsets the increased design effect and power does not change appreciably with size of enrolled communities when transmission is moderate (Fig. 4F).

### Analysis methods for a cRCT

In answering our primary research questions, we explored a range of analysis methods for the cRCT. We found that a Cox PH model with Gaussian-distributed frailty had significantly elevated Type I error (see online <u>Shiny App</u>). Fortunately, two common approaches to analyzing clustered survival data, a Cox PH model with gamma-distributed frailty or robust standard error estimation, were the best methods in terms of power and validity. The robust SE analysis has higher power than the gamma-frailty model when transmission is low. However, the model doesn’t account for heterogeneity in hazard rates in its estimate of the vaccine effect, leading to a downward bias that is particularly apparent when R_0_ is high, as seen in Figure 2. The gamma-frailty model is not susceptible to this bias.

## DISCUSSION

Traditional comparisons of cRCTs versus iRCTs that focus on within-cluster correlation and the design effect should also consider other ways in which the unit of randomization affects RCT power. Although an iRCT and a cRCT answer different research questions (measuring direct and total effects, respectively), a positive finding for either could arguably lead to the same policy outcome, especially during an epidemic (15). For example, rVSV-EBOV was approved for use in the DRC in 2017 based on the findings of *Ebola, ça suffit!*, a cRCT (16). We show that a cRCT’s ability to measure both indirect and direct effects can partially compensate for the loss of power due to clustering. Theoretical calculations suggest that cRCTs may exhibit greater statistical efficiency than iRCTs in some low R_0_ scenarios. However, simulations that more realistically capture stochasticity in transmission suggest that iRCTs remain more powerful than cRCTs conducted in the same trial population. In low transmission settings the difference in power between them may be small, although for R_0_ values lower than considered here a risk-prioritized design (such as ring vaccination) would be preferable, and these results should be examined separately in this context.

The above comparisons between cRCTs and iRCTs can be extended to examine cRCTs of different cluster sizes (which is particularly apparent once noting an iRCT can be considered a cRCT with cluster size of one). For instance, within cRCT designs, enrolling more individuals from the same cluster is generally less statistically efficient than enrolling individuals in a new cluster. Previous work has argued that the ICC often decreases with cluster size, mitigating some loss of efficiency with larger clusters (10), and demonstrated how cross-contamination may increase when cRCTs are run in clusters of fewer individuals, reducing the effect to be estimated and thus power. Cross-contamination occurs either via transmission between intervention and control clusters or inadvertent receipt of intervention by control clusters, both of which are less likely when clusters are separated in space (17), or are sufficiently large that they are less impacted by external populations (10).

Here, we show that, even in the absence of cross-contamination, indirect effects in themselves can mitigate the loss of efficiency caused by the increasing design effect associated with fewer, larger clusters. To our knowledge, this fact has been alluded to but the effect on power has never been quantified (18, 19). Another counter-intuitive finding arises from the fact that, because larger communities experience a greater influx of transmission imported from elsewhere, enrolling fewer but larger communities may yield a greater attack rate, and thereby partly or fully compensate for the loss in power due to the design effect. This result is dependent on the relationship between case importation rate and community size. Consequently, this will differ by disease and population setting and may only be true in scenarios when a pathogen is not endemic to trial communities and the probability of pathogen introduction into a community is relatively low.

Our findings highlighted the importance of adequately accounting for heterogeneity between study clusters while maintaining the nominal false positive rate and maximizing power. We limited the methods to those widely used and found that a Cox PH model with gamma-distributed frailty performs best overall; although when R_0_ is low, a Cox PH model with robust SE may be superior.

The results presented here are part of a body of work demonstrating the utility of simulation when considering the design of vaccine trials for infectious diseases (7). It is only by including transmission dynamics in models that we are able to quantify the relative strength of clustering and indirect protection in affecting trial power. Our study is intended to explore these effects more generally, but we expect our findings to be relevant to investigators considering cRCT design, whether or not they develop a full-fledged trial simulation study during the planning phase. Theoretical work on trial design can help prepare stakeholders to rapidly design trials in the face of unexpected epidemics of emerging pathogens. However, it is important to note that sample size is only one of many factors that must be taken into consideration when planning a vaccine trial. Considerations of logistics, cost, ethics, acceptability or the particular research question of interest may, in certain contexts, hold priority.

There are at least two sources of intracluster correlation in a cRCT for an infectious disease: transmission between individuals within a cluster, and the shared characteristics of individuals within a cluster. When R_0_ is large enough, any outbreak that takes off will infect many individuals in a community so all clusters either have attack rate close to 0% or 100%. In such cases, there is very little within-cluster variance and the total variance comprises chiefly between-cluster variance, leading to ICCs approaching 1. Clustering due to shared characteristics can arise for many reasons, e.g. within-community similarities in behavior, health, or proximity to source populations. Intracluster correlation, whether due to transmission or to shared characteristics in clusters, increases the design effect. Given these different sources of clustering, and the fact that we observed ICCs ranging from 0.05 to 0.8 in our simulations, it is critically important that ICCs are reported by study investigators when presenting the results of a cRCT as this may aid in planning for future trials (20).

Our analysis neglects some aspects of a realistic population in which a trial is conducted. For example, we do not consider the second source of clustering described above (i.e. shared characteristics). More broadly, modeled individuals do not vary in characteristics other than degree and the community to which they belong, whereas real populations would vary in age structure, proximity to the epicenter of the epidemic, and other variables that would predict disease incidence. By ignoring these characteristics we underestimate the extent of clustering in a cRCT and overstate its power. This makes more robust our conclusion that the iRCT is always more powerful than the cRCT in the situations considered.

We have conceptualized the population structure as being a number of small groups separated in space so that there is minimal transmission between communities; in reality, population structure is likely to be less distinct. We have not considered permanent or temporary migration, nor secondary structure within communities (i.e. households). Moreover, real-life degree distributions have a heavier tail (due to superspreading (21)) than considered here; though a sensitivity analysis shows our results are robust to this assumption (see online <u>Shiny App</u>).

We find that the general principle that enrollment of fewer, larger clusters leads to decreased power is strongly dependent on the relationship between community size and rate of importation. Our base assumption that importation frequency proportional to the square root of community size is based on a finding for measles (11). For other diseases the community-level importation rate may be independent of community size, in which case the increased design effect would entirely dictate the loss of power as community size increases. Our conclusions should thus be considered in the context of each specific disease and population.

The indirect effect of vaccination should be considered along with clustering in calculating the power of a cluster-randomized trial and in comparing different trial designs for interventions against infectious diseases. Using simulation we show that it does not always increase power to enroll more, smaller clusters into a cRCT, when doing so is associated with reduced indirect protection to vaccinated individuals or importation of infection into the study population. Still, while cRCTs measure a greater vaccine effect than iRCTs, we found that iRCTs are generally more powerful, though their power may be comparable in low-transmission settings.

## List of supplementary material

Supplementary Materials and Methods.

**Figure S1.**
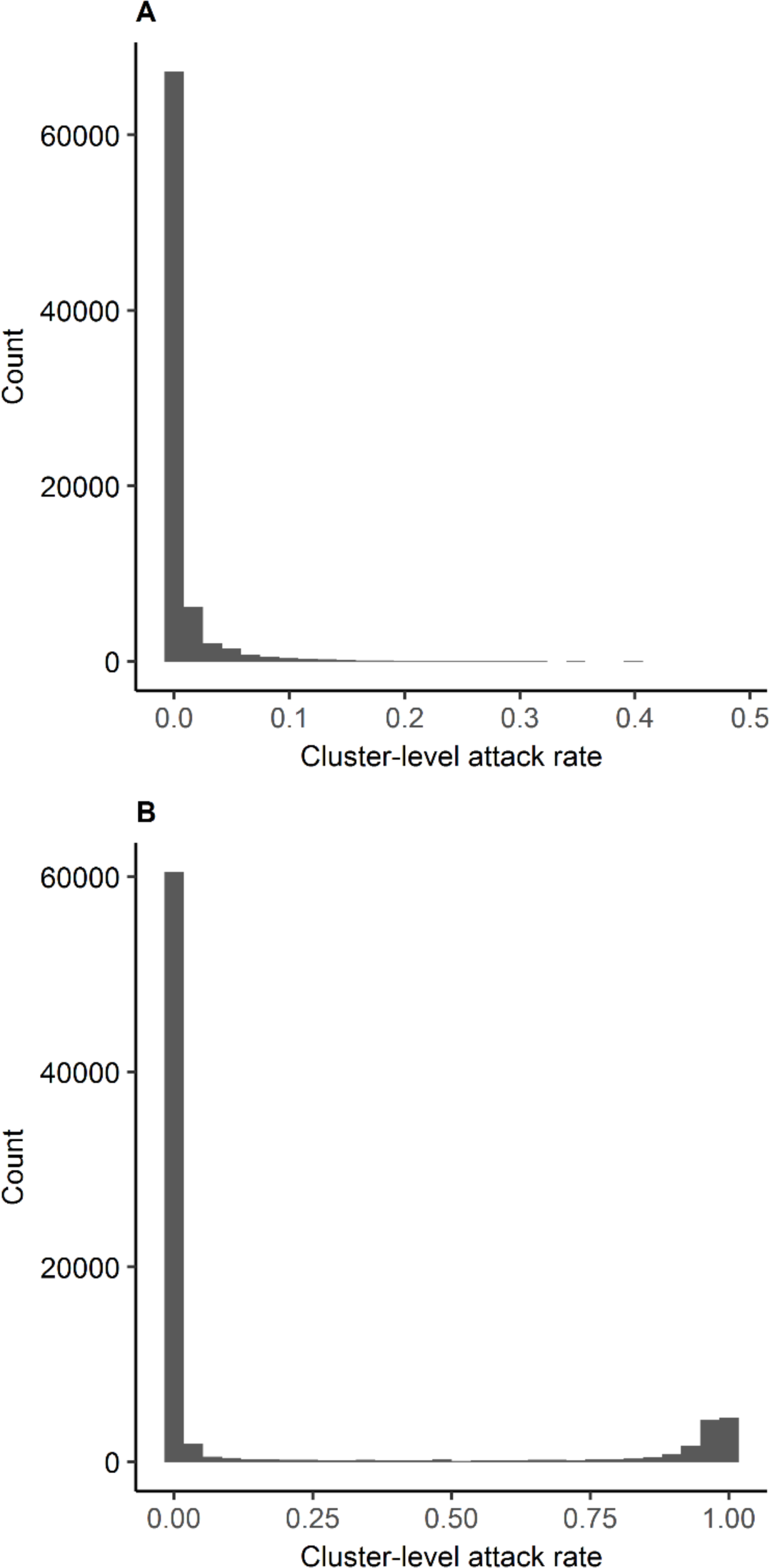
Relationship between R_0_ and distribution of cluster-level attack rates. Histogram of cluster-level attack rate for R_0_=0.6 (**A**) and R_0_=3 (**B**).

**Figure S2.**
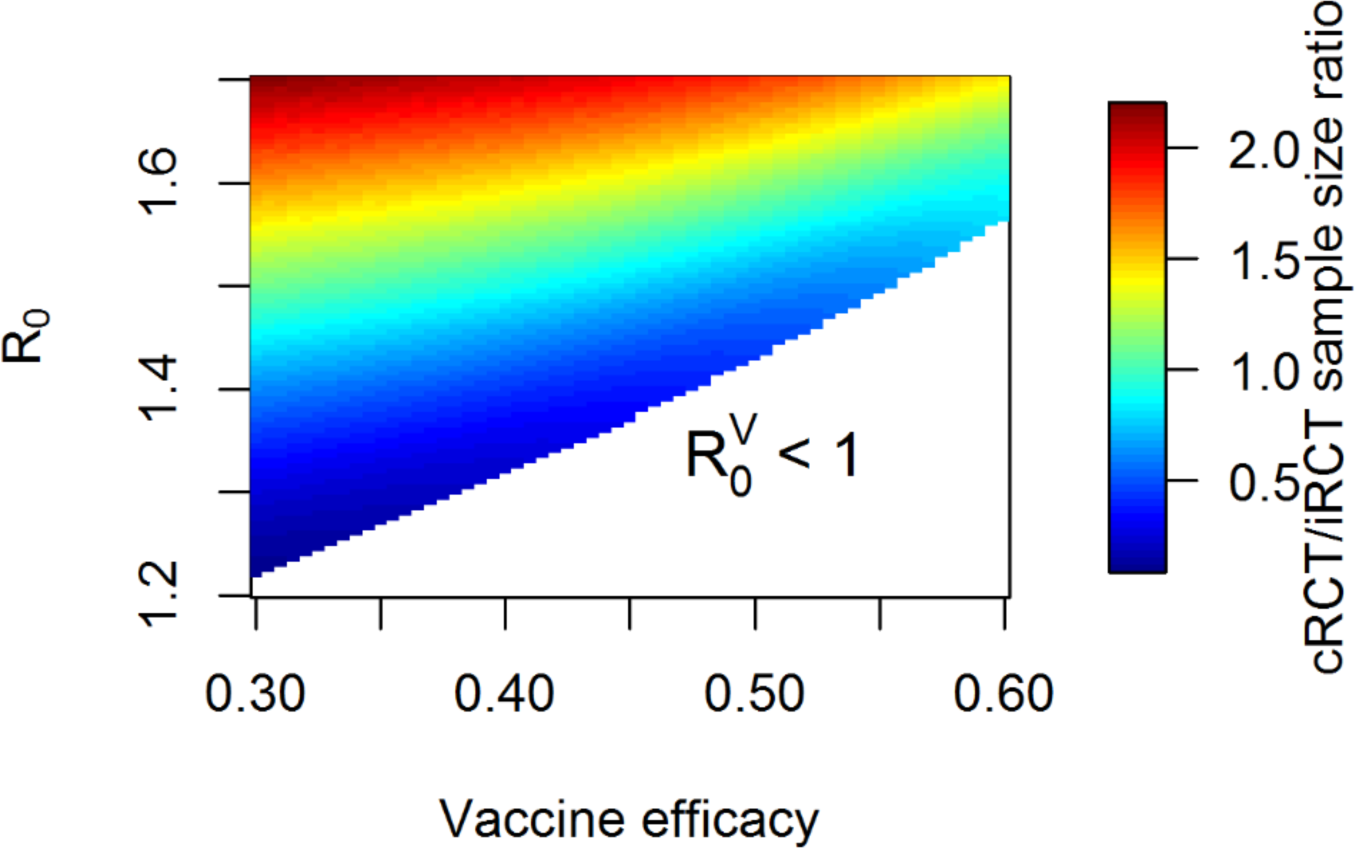
cRCT/iRCT necessary sample size ratio by R_0_ and vaccine efficacy from theoretical model. Ratio of necessary sample size for 90% power to detect vaccine effect for a cRCT (total effects) relative to an iRCT (direct effect) with a hazard rate-based analysis, varying R_0_ and true vaccine efficacy. Final size equations apply only when 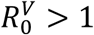.

## Supplementary Materials

### Methods

#### 1. Theoretical model

Our initial theoretical analysis is based on final size and outbreak probability calculations rather than simulation, but otherwise the trial population has the same structure as in the simulation model. In particular, we assume that a proportion of communities receive a single disease importation, and any outbreak that arises from an importation runs until there are no longer any infectious individuals.

For a community in which there are no vaccinees, the standard final size equation (1) applies for the cumulative incidence, namely CI solves **CI = 1 − e^-R_0_CI^**, when R_0_>1. Similarly, the proportion of communities with importations in which an outbreak will occur, x, solves the same equation. For a community in which a proportion *p* of the individuals are vaccinated with vaccine efficacy VE, the equations for the CI among the vaccinated and unvaccinated, CI_V_ and CI_U_ respectively, are

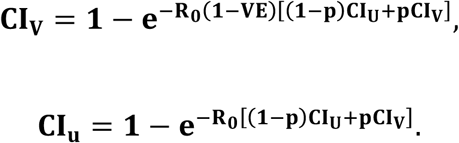

The outbreak probability in a community in which a proportion *p* of the individuals are vaccinated, *x_V_*, solves the equation 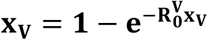, where 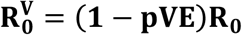. Sample size calculations were based on a hazard rate analysis, with vaccine effect estimated in both trial designs as 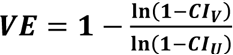(2). Specifically, number of individuals needed to achieve 90% power to detect vaccine effect was given by 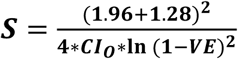(3), where *CI_O_* is the cumulative incidence of infection in the trial population. For the cRCT, this sample size is multiplied by the design effect as defined in the main text, with ICC calculated using the ANOVA method (4). We calculate necessary sample size to achieve 90% power for an iRCT (in which half of the vaccinees in each study cluster are vaccinated) and for a cRCT (in which half of the study clusters have all participants vaccinated, and the other half are given control), and plot the ratio of the necessary sample size for a cRCT compared to an iRCT. Areas of parameter space in which this ratio is less than 1 are indicative of parameters for which the cRCT is theoretically more efficient at detecting the total effect than the iRCT is at detecting the direct effect.

When R_0_<1, the size of an outbreak in a large population is given by 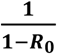 but this formula does not apply to small communities, especially when R_0_ is close to 1. Therefore, we restrict the theoretical analyses to parameter combinations when R0 in vaccinated communities in the cRCT is greater than 1, assuming that any qualitative results we saw in this parameter space would be maintained as R_0_ crosses 1.

#### 2. Simulation

The main population model is a standard deterministic susceptible-exposed-infectious-removed (SEIR) compartmental model, with three exposed and three infectious compartments to yield gamma-distributed incubation and infectious periods. We assumed a time-varying transmission rate in the main population, so that the importation rate into the communities is proportional to the prevalence of infection in the main population, and disease natural history parameters representative of the 2014-2015 Ebola epidemic in Liberia (5).

The disease model in the communities is a stochastic susceptible-exposed-infectious-removed (SEIR) model. Each susceptible individual has a daily hazard of becoming infected and moving into the exposed compartment from two sources: the daily hazard of infection from each infectious *neighbor* is *β*, and the daily hazard of infection for an individual in community *i* from the main population is *F_i_I*, where *I* is the prevalence of infectious individuals in the main population and *F_i_* is a proportionality constant reflecting the degree of contact between the main population and the *i*^th^ community.

The hazard rate of introduction into the study population is time-varying with the progression of the epidemic in the main population, and we calibrate the constant of proportionality in each cluster F_i_ using an assumed rate of importation events, *M_i_* cases/year. The formula that connects these two quantities is 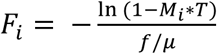, where *f* is the final size of the epidemic in the main population, *μ* is the mean infectious period, and *T* is the length of the epidemic in years. We model the relationship between importation rate and community size in two ways. Firstly, for community *i* we assume 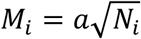, where *N_i_* is the community size (6), and the *per capita* importation rate in community *i* is 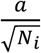, where the constant *a* determines the magnitude of the importation rate. Secondly, we assume *M_i_* = *a′N_i_*, so that the *per capita* importation rate in community *i* is *a’*. The values for *a* and *a’* were chosen so that a community of size N_i_=100 had on average between 0.25 and 1 introductions over the course of a two-year epidemic.

The transmission rate *β* in the main population varied with time using the formula 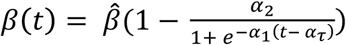. Parameters were chosen to give a reasonable fit to weekly Ebola incidence data from Liberia. Specifically, 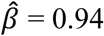, *α_1_* = 0.19, *α_2_* = 0.6, *α_τ_* = 27.79. The average incubation/latent period is 7.14 days and the average infectious period is 3 days.

We assume that the incubation and latent periods are concurrent, meaning that symptom onset occurs when infectiousness begins. Once infected, individuals spend a number of days in the exposed compartment drawn from a gamma distribution with mean 9.7 days and SD 5.5 days before moving into the infectious compartment (7). They spend a number of days in the infectious compartment drawn from an independent gamma distribution with mean 5 and SD 4.7 based on data on the time to hospitalization (7), after which they move into the removed compartment. For simplicity and to generalize away from the Ebola epidemic, we assume no *post mortem* transmission, meaning that whether an individual dies or recovers does not affect the estimated efficacy or power of the trial.

Once enrolled, individuals are followed for a number of days and, for infected individuals, time from enrollment to symptom onset is recorded. Individuals who never develop symptoms are censored at the end of the study; there are no other sources of censoring. The vaccine is multiplicative leaky (8), reducing susceptibility to infection by a factor (1-VE) and having no effect on those who are already exposed or infectious when vaccinated, and no effect on the progression or infectiousness of vaccinated individuals who become infected. We assume the protective efficacy of the vaccine starts on the day of vaccination.

